# Whole methylome changes in Barrett’s metaplasia, dysplasia, and cancer

**DOI:** 10.64898/2026.10.30.763255

**Authors:** Yi Jin Liew, Thejaani P. Udumanne, Kyle Catalan, Amali Ariyavidana, Ka Ki Michelle Lee-Ng, Dana Pascovici, Tao Yang, Priyanthi Kumarasinghe, Duncan McLeod, Ian Brown, Michael J. Bourke, Sarah J. Lord, Reginald V. Lord, Jason Ross

## Abstract

The clinical management of Barrett’s esophagus (BE) and the risk of esophageal adenocarcinoma (EAC) is dependent on the presence or absence of dysplasia. The histopathological diagnosis of dysplasia is unreliable however, and would benefit from additional diagnostic criteria. Here, we obtained EM-seq whole-genome methylation profiles from 86 samples across the BE/EAC disease spectrum, notably supplementing the scant existing methylation evidence for low-grade and high-grade dysplasia. While the bulk of methylation changes occur in the transition from normal squamous epithelium to intestinal metaplasia (non-dysplastic Barrett’s esophagus), we identified a smaller set of changes that mark the onset of dysplasia and progression to EAC. Methylation changes were broadly independent of copy number variation. They were tightly anticorrelated with RNA-seq expression for some genes. Deconvolution of the bulk methylation signatures into constituent cell types revealed four sample groupings enriched with a particular cell type (squamous, columnar, lymphoid and erythroid cells), somewhat reflecting existing clinical classifications. We outline potential markers that could improve the diagnosis of metaplasia and dysplasia in EAC.

## Introduction

Esophageal adenocarcinoma (EAC, includes adenocarcinoma of the gastroesophageal junction) and esophageal squamous cell carcinoma are the two major subtypes of esophageal cancer, which ranks as the 11^th^ most prevalent cancer and 7^th^ leading cause of cancer mortality globally (YANG *et al*. 2024). EAC is the predominant subtype in Western countries, where its incidence in some areas has increased as much as 7-fold since 1973 (HANG *et al*. 2022). EAC has one of the highest case-fatality ratios of all cancers, with a ∼20% five-year survival rate (SIEGEL *et al*. 2025).

The main risk factor for EAC is the presence of Barrett’s esophagus (BE), in which the normal squamous lining of the distal esophagus is replaced with a metaplastic columnar cell epithelium with mucous-secreting goblet cells. BE occurs in response to gastroesophageal reflux disease (GERD) and is estimated to occur in 3% of adults in Western and 1% in non-Western countries (QUMSEYA *et al*. 2019). The development of EAC in most patients is thought to occur via a multistage carcinogenesis process in which the presence of dysplasia in BE (low-grade dysplasia, LGD; or high-grade dysplasia, HGD) is associated with an increased cancer risk and guides patient management (KILLCOYNE & FITZGERALD 2021).

Unfortunately, histopathological examination for the diagnosis of dysplasia is unreliable (MONTGOMERY *et al*. 2001; SONWALKAR *et al*. 2010), especially for LGD; variability in the pathological diagnosis of LGD is reflected in the very wide (∼1% to >10%) range in estimates of progression to HGD/EAC in patients with LGD (SINGH *et al*. 2014; MOOLE *et al*. 2016). Adding accurate molecular information to the histopathological criteria might improve this situation, which might also help identify those patients for whom treatment with endoscopic therapy, rather than surveillance alone, is appropriate.

Previous studies have identified widespread methylation changes associated with non-dysplastic BE (NDBE) and EAC (XU *et al*. 2013; KRAUSE *et al*. 2016; YU *et al*. 2019; JAMMULA *et al*. 2020; NAEINI *et al*. 2023) by assaying a subset of methylated cytosines (< 1M CpGs) in the human genome with commercial Infinium BeadChip methylation arrays; BE with dysplasia tissues were only assayed in earlier non-array studies e.g., Eads et al. (2001). As EAC is a cancer with high mutational burden and somatic copy number variation (KILLCOYNE & FITZGERALD 2021), the observed methylation changes could potentially correlate with genomic instability (MARTIN-TRUJILLO *et al*. 2017; GARG *et al*. 2021), with implications for the design of universal markers for the disease. In EAC, early evidence suggests significant methylation changes are largely independent from copy number variation (NAEINI *et al*. 2023), but this evidence is limited to the subset of assayed cytosines.

In this study, we analyzed whole methylome data with enzymatic methylation sequencing (EM-seq; > 25M CpGs) from 86 patients, including patients with LGD and HGD, who were enrolled in a prospective study (NA *et al*. 2021). We explore genome-wide methylation changes across the normal squamous (NSq)-to-EAC disease progression spectrum; we determine whether copy number variation confounds the identification of differential methylation; and we investigate whether the deconvolution of bulk methylome data into constituent tissues may provide insights into the biology of disease progression and aid biomarker discovery.

## Methods

### Patient cohort, specimen collection and disease classification

Ethics approval was obtained through the Progression of Barrett’s Oesophagus to Cancer Network (PROBE-NET) study (2019/ETH03452). All participants provided written informed consent prior to recruitment and sample collection. Descriptive cohort characteristics were generated with the “furniture” (v1.11.0) R package (BARRETT & BRIGNONE 2017). Significance testing were chi-square tests (package defaults for categorical data).

For patients with BE, tissue biopsy samples were collected as per Seattle protocol (1 cm above GEJ, and then every 2 cm proximally within the columnar lining). Normal squamous mucosa biopsies were taken from ∼20 cm from incisors. For EAC patients, the tissue samples were collected from within the cancer, and from adjacent BE when present. The tissue samples were immediately placed in RNAlater solution (Thermo Fisher Scientific, cat. #AM7020) for stabilization and stored at 4 °C for 12–24 hours to ensure tissue preservation, then transferred to –80 °C for long-term storage. Central sections of each specimen were formalin-fixed, paraffin-embedded and processed for H&E staining and subsequent histopathological assessment. These H&E sections were reviewed by at least two expert Barrett’s esophagus pathologists who had to agree to confirm the diagnosis.

We additionally classified samples as low-or high-risk according to whether worse pathology was present in the esophagus at the time of endoscopy. A patient with high-risk LGD could, for example, have HGD or EAC in other biopsies from the same endoscopy, whereas a patient with low-risk LGD had LGD as the worst pathology diagnosed at that endoscopy. Risk was not assigned to EAC samples. Detailed cohort characteristics are provided in Supplementary Table S1.

### DNA extraction and library preparation from selected samples

We selected *n*=86 fresh-frozen esophageal tissue biopsies for DNA extraction and library preparation (10 NSq, 33 NDBE, 22 LGD, 19 HGD, 13 EAC) from 72 patients. DNA was extracted with the QIAamp Fast DNA Tissue Kit (Qiagen cat no. 51404, lot no. 172015357) following manufacturer’s recommendations with a few optimizations. We increased the proteinase K digestion time from 10 minutes to 30 minutes; performed an extra washing step with buffer AW2 (i.e., two AW2 washes in total); and for the elution step, after adding in DNase-/RNase-free water onto the filter, the columns were incubated at room temperature for 4 minutes prior to the final centrifuge spin.

High molecular weight unsheared DNA (> 500 ng) from 86 samples were submitted to the Australian Genome Research Facility (AGRF, Melbourne, Australia), following AGRF’s service guidelines for the Whole Genome Enzymatic Methylation Sequencing (WGEMS) service. EM-seq libraries were prepared from the submitted samples and two spike-in controls (unmethylated lambda and fully methylated pUC19), then sequenced on a NovaSeq 6000 (Illumina).

### Processing raw reads for CpG methylation levels

We used https://github.com/lyijin/bismsmark, a thin Snakemake (MOLDER *et al*. 2021) wrapper around the Bismark pipeline (KRUEGER & ANDREWS 2011) that facilitated its deployment on the CSIRO High Performance Computing clusters, to process raw reads into tab-delimited files containing per-position methylation levels. In brief, raw reads were trimmed 10 bp on every end according to Bismark’s best practices, then mapped against the GRCh38.p13 genome with lambda and pUC19 sequences (Supplementary Table S2). Methylation levels were then called from paired-end reads mapped in the correct orientation with a maximum insert size of 1,000 bp (the default of 500 bp was more appropriate for WGBS libraries that tend to be smaller than EM-seq libraries). Per-CpG coverages were combined across strands with Bismark’s coverage2cytosine tool. We retained CpGs with any coverage in > 95% samples and with the lower quartile of coverage values > 5 across all samples.

### Identification and functional analysis of differentially methylated regions

The identification of differentially methylated regions (DMRs) was done using metilene (JUHLING *et al*. 2016). DMRs were filtered on two additional criteria: minimum change in methylation level over baseline (β) > 0.1 and qval < 0.01. Genomic features of methylated CpGs were added with self-written R scripts, focusing on whether they resided in CpG islands, CpG shores or neither; whether they were in promoters, gene bodies, or intergenic regions; whether they were in repeat elements; and the name of the closest protein-coding gene. With these additional annotations, clusterProfiler v4.19.7 (WU *et al*. 2021) was used to identify GO and KEGG pathways enriched with genes containing hypermethylated or hypomethylated DMRs (analyzed separately). Code and intermediate files for these analyses are in our GitHub repository, under subfolders numbered 8–11.

### Identifying copy number variation in EM-seq data

CNVkit v0.9.13 (TALEVICH *et al*. 2016) was used to identify genomic regions with copy number variation (CNV) in non-NSq samples (as the 10 NSq samples were used to form the “normal” copy number baseline). NSq samples, by definition, had no CNV; they were not included in subsequent analysis. The most notable non-standard parameter used was the size of the genomic windows, which we set at 20 kb (--target-avg-size 20000), higher than the loose consensus (for WGS data) at 5 kb. After testing, we opted for larger buckets given some of our samples had poor coverage.

Despite the use of larger buckets, samples with the poorest coverages still resulted in noisy CNV calls (which results in a “striping” pattern in heatmaps due to false positive amplifications/deletions). For CNV analyses, we retained 54 (of 76 non-NSq samples) with overall coverage > 12. Exact commands to replicate our analyses can be found in our GitHub repository, under the subfolder numbered 12.

### Correlation with expression patterns

Expression data from the same samples, where available, was obtained from (UDUMANNE *et al*. 2026); both studies were consistent in having performed pairwise comparisons across the NDBE-to-EAC spectrum. An R script was written to search for differentially methylated promoters that were co-located in a differentially expressed gene, tabulating the direction and magnitude of promoter methylation and gene expression. Quadrant plots were generated with ggpp v0.6.0.

To investigate the direct correlation between methylation and expression, we used the subset of samples with both whole methylome and transcriptome data (*n* = 46; Fig. 1A). Pearson correlations and *p* values were computed in R for individual genes, followed by Benjamini-Hochberg correction on the *p* values to control for multiple testing. Significance was set at post-corrected *p* < 0.01. Exact commands to replicate this section are in the subfolder numbered 13.

**Fig. 1.**
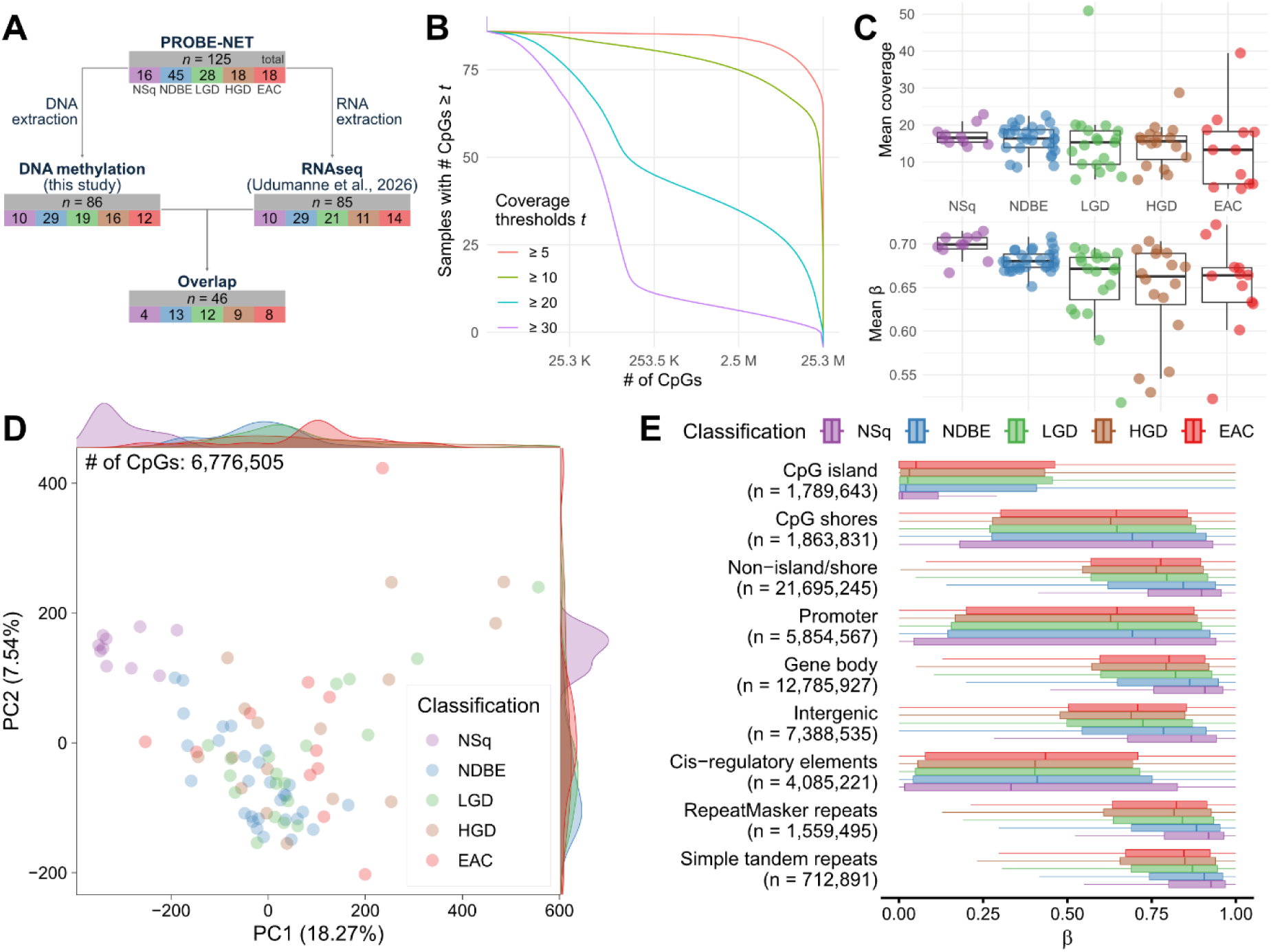
Overall methylation patterns across the five stages of BE/EAC progression. **(A)** Study set-up, with overall number of samples (in gray) and individual stages across the BE/EAC progression spectrum. Whole-genome methylation data for *n* = 86 samples were obtained for this study, of which a subset of *n* = 46 also has expression data. **(B)** Coverage plot across all samples post-filtering of low-coverage CpGs (25.3M positions). **(C)** Mean coverage (top panel) and mean beta (bottom panel) for each sample, binned by disease classification. While the poorest covered samples tend to be EAC, coverage differences across any group pairings were not statistically significant. **(D)** PCA across all 86 samples to illustrate overall similarities of methylation patterns. **(E)** Mean methylation levels of specific genomic contexts, separated by disease classification. Colors and abbreviations are standardized across subplots. NSq: normal squamous, purple; NDBE: non-dysplastic Barrett’s esophagus, blue; LGD: low-grade dysplasia, green; HGD: high-grade dysplasia, brown; EAC: esophageal adenocarcinoma, red.

### Deconvolution of bulk methylation profiles into tissue-specific compositions

Bulk methylation profiles were deconvoluted with CelFiE-ISH (UNTERMAN *et al*. 2024) with the cell-type-specific methylation atlas (LOYFER *et al*. 2023). Both resources were based on the GRCh37 (hg19) human genome; but our work so far had revolved around the GRCh38 (hg38) genome. We noticed several oddities when we converted GRCh37 coordinates to GRCh38 with liftOver (HINRICHS *et al*. 2006): some coordinates failed to convert; while some were lifted into a completely different context. Following information gleaned from the Supplementary section of Loyfer et al. (2023), we replicated the creation of the methylation atlas on GRCh38 coordinates. Exact commands used to replicate our efforts (and timesavers for others, e.g., the GRCh38-equivalent methylation atlas file) can be found in our GitHub repository, under the subfolder numbered 14. All samples were deconvoluted with the GRCh38-equivalent pipeline to produce a table with per-sample, per-tissue abundance fractions.

Unsupervised hierarchical clustering of these values to produce cell-type clusters was carried out in R (subfolder 14). Based on these clusters, promoter methylation (subfolder 15) and gene expression patterns (subfolder 16) were further analyzed in R.

## Results

### Whole-genome methylome sequencing with EM-seq

Data was obtained from *n* = 86 samples (10 NSq, 29 NDBE, 19 LGD, 16 HGD, 12 EAC; Fig. 1A, Table 1, Supplementary Table S1). We used EM-seq instead of whole genome bisulfite sequencing (WGBS) as the enzymatic conversion of DNA is less destructive, resulting in more even coverage across the genome and more accurate methylation level readouts from GC-rich loci (VAISVILA *et al*. 2021; GUANZON *et al*. 2024). CpGs with consistently low coverages across all samples were removed, retaining 25,349,327 CpGs for further analysis (Fig. 1B). Overall coverage was independent of disease classification (*p* = 0.73, single-factor ANOVA; Fig. 1C).

**Table 1.** Descriptive details for normal, Barrett’s, and EAC tissues. All EAC samples are high-risk by definition (see Methods).

|  | **Classification** |  |  |  |  |  |
|---|---|---|---|---|---|---|
|  | **NSq** | **NDBE** | **LGD** | **HGD** | **EAC** | ***p* value** |
|  | *n* = 10 | *n* = 29 | *n* = 19 | *n* = 16 | *n* = 12 |  |
| **Sex** |  |  |  |  |  | 0.29 |
| M | 8 (80%) | 27 (93%) | 18 (95%) | 12 (75%) | 11 (92%) |  |
| F | 2 (20%) | 2 (7%) | 1 (5%) | 4 (25%) | 1 (8%) |  |
| **Age** |  |  |  |  |  | 0.044 |
| Mean (SD) | 61.3 (12.5) | 59.7 (10.9) | 68.1 (8.2) | 66.8 (12.0) | 67.9 (11.7) |  |
| **Risk** |  |  |  |  |  | 0.01 |
| L | 5 (50%) | 17 (59%) | 6 (32%) | 6 (38%) | - |  |
| H | 5 (50%) | 12 (41%) | 13 (68%) | 10 (62%) | 12 (N/A) |  |

### Overall methylation patterns mainly depend on cell type

To visualize the overall methylation patterns across the five pathology stages, we carried out a principal component analysis (PCA) on ∼50% of all positions with sufficient coverage across all samples. The 10 NSq samples clustered closely together, but the remaining 76 non-NSq samples did not cluster as tightly by disease classification (Fig. 1D; Supplementary Fig. 1).

There was a global hypomethylation across the BE/EAC stages, with the steepest decline at the transition from NSq to intestinal metaplasia (NDBE, Fig. 1C). Despite this global hypomethylation, we observed focal hypermethylation occurring in CpG islands when genomic contexts were averaged separately (Fig. 1E). For other genomic contexts, in general, methylation levels trended away from the extremes (fully unmethylated/methylated) towards the midpoint in worse pathology tissues (Fig. 1E).

**Supplementary Fig. 1.**
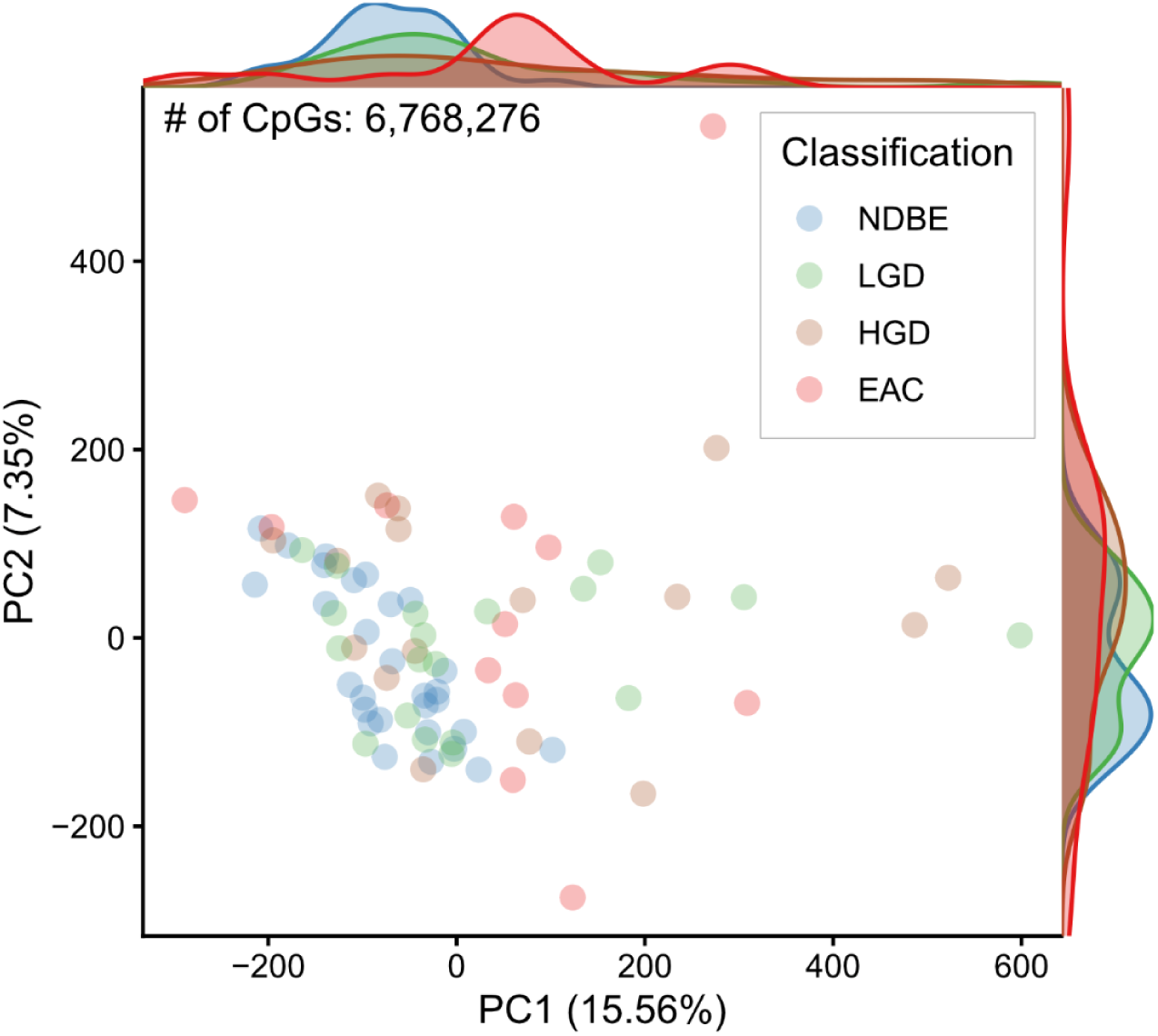
Principal component analysis on non-NSq samples to discern higher-order patterns in Barrett’s esophagus. While NDBE samples have methylation patterns that were most consistent (i.e., least variable), there is no universal change in methylation patterns that cleanly separates NDBE from LGD or HGD. Repeating the principal component analyses on smaller subsets (e.g., excluding EAC samples, focusing on samples with higher sequencing coverages, analyzing the top 100,000 most variable methylation positions) did not result in any materially different plots (exploratory plots available on GitHub). Samples are colored by stage (NDBE: blue; LGD: green; HGD: brown; EAC: red).

### Bulk of hypermethylation driven by intestinal metaplasia (NDBE)

We used metilene (JUHLING *et al*. 2016) to identify differentially methylated regions (DMRs) across six biologically relevant pairwise comparisons (Fig. 2A; Supplementary Tables S3–14). Four comparisons tracked BE/EAC progression steps, i.e., the transition to metaplasia (NDBE vs. NSq); onset of dysplasia (LGD vs. NDBE); worsening of dysplasia (HGD vs. LGD); and development of invasive carcinogenesis (EAC vs. HGD). Two other comparisons with potential clinical utility in addressing current diagnostic gaps were included (HGD/EAC vs. NDBE; and EAC vs. NSq).

**Fig. 2.**
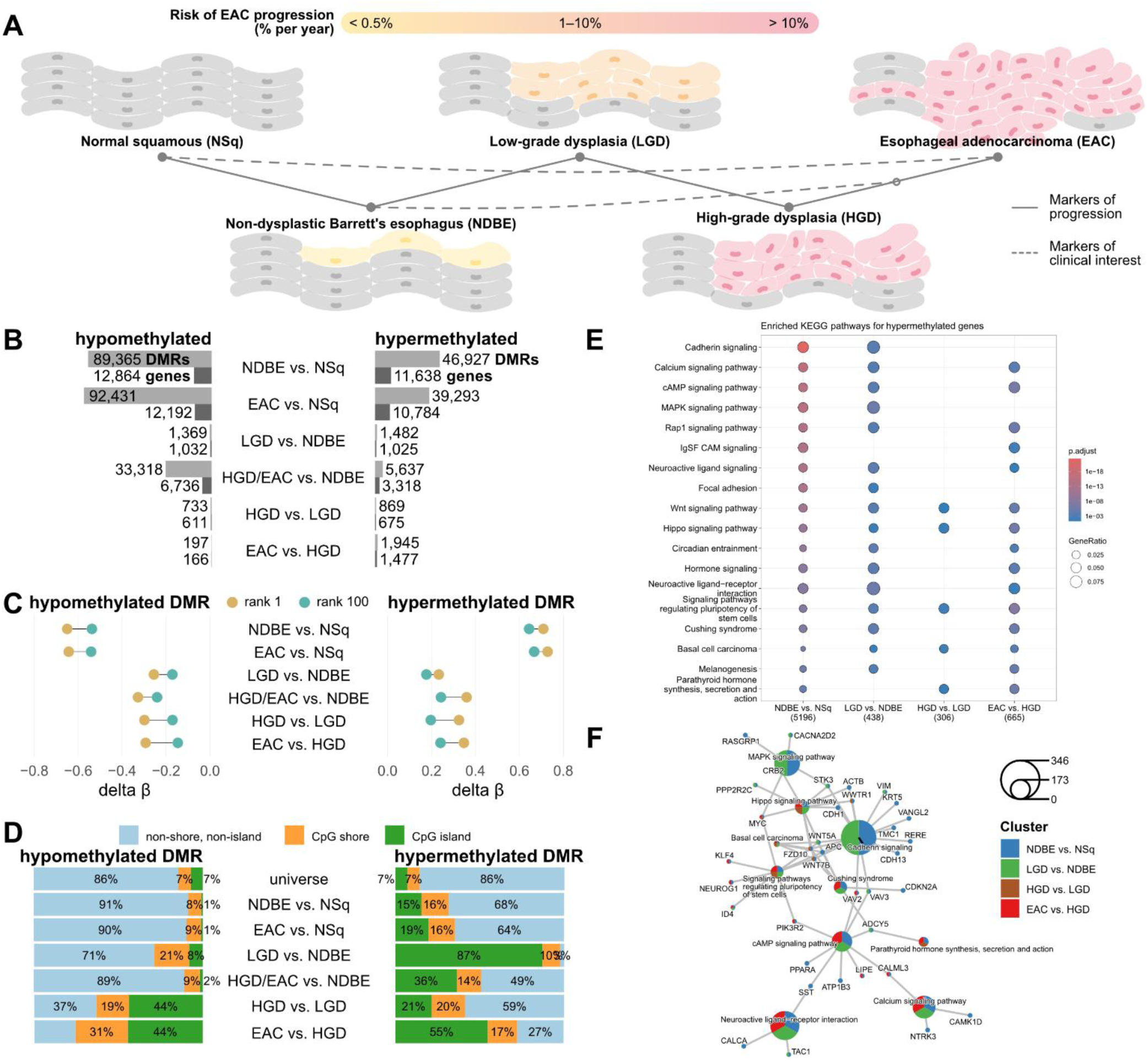
Overview of DMRs identified from whole-methylome data. **(A)** A sketch of the progression of EAC, annotated with the six pairwise comparisons carried out in this study (solid/broken lines). In all subsequent subplots, all pairwise comparisons are labelled following the “more vs. less severe pathology” pattern, i.e., hypermethylated DMRs for LGD vs. NDBE have higher regional methylation levels in LGD than in NDBE. **(B)** Tallies of differentially methylated regions (DMRs, light gray) and genes (dark gray) across six comparisons of interest. **(C)** Dumbbell plot showing the extent of methylation level changes of the DMRs (delta beta; rank 1 in brown, rank 100 in green). **(D)** Genomic contexts of DMRs, classified by CpG island (green), CpG shores (orange), or neither (blue). The universe refers to the genomic context of all CpG dinucleotides with valid data. **(E)** KEGG pathways enriched with hypermethylated genes across sequential EAC progression. Dysregulated pathways could occur across multiple pairwise comparisons (denoted by circles). Colors (red/blue) indicate *p* value significance; circle sizes indicate the fraction of genes in that pathway that were hypermethylated. **(F)** Gene networks highlighting the presence of known EAC and top 50 most hypermethylated genes. Genes were colored by stages where they exhibit hypermethylation: at transition to metaplasia (NDBE vs. NSq, blue), at onset of dysplasia (LGD vs. NDBE, green), worsening dysplasia (HGD vs. LGD, brown) and at carcinogenesis (EAC vs. HGD, red).

While there were more hypomethylated DMRs at the early stages of disease, a reflection of the initial marked global hypomethylation in NDBE, hypermethylated DMRs exceeded hypomethylated ones with progression of dysplasia to cancer (Fig. 2B). The magnitude of differential methylation (delta beta) was consistently stronger for hypermethylated DMRs (Fig. 2C). Hypermethylated DMRs were disproportionately enriched in CpG islands; while hypomethylated DMRs tended to be outside of CpG islands or shores—excepting comparisons involving HGD (Fig. 2D). Functional analysis of the DMRs revealed that hypermethylated DMRs were more likely to be associated with genes in pathways that modulate cell growth, cell-cell adhesion, and included two annotated cancers in the KEGG database: gastric cancer and basal cell carcinoma (Fig. 2E, 2F).

To investigate whether the risk of sampling error (in which the sampled biopsies do not include the worst pathology present) could be mitigated, we were interested in understanding whether methylation changes in biopsies with more benign pathology could predict the presence of worse adjacent pathology. Our samples were therefore divided into “high-risk” (sampled tissues were adjacent to worse pathology) and “low-risk” (worse pathology was not present in adjacent biopsies). Whilst risk-based DMRs are generally less frequent than those from inter-stage comparisons (Supplementary Fig. 2A; Supplementary Tables 15–22), risk-based DMRs from LGD and HGD are about as frequent as those denoting the onset of dysplasia (LGD vs. NDBE) and worsening of dysplasia (HGD vs. NDBE; Supplementary Fig. 2A). DMRs also tended to be hypermethylated, and similarly disproportionately located in CpG islands (Supplementary Figs. 2B, 2C). In terms of enriched functional pathways, “transcriptional misregulation in cancer” is unique to risk-based DMRs, suggesting the presence of risk-based clues in other –omic layers (Supplementary Fig. 2D).

**Supplementary Fig. 2.**
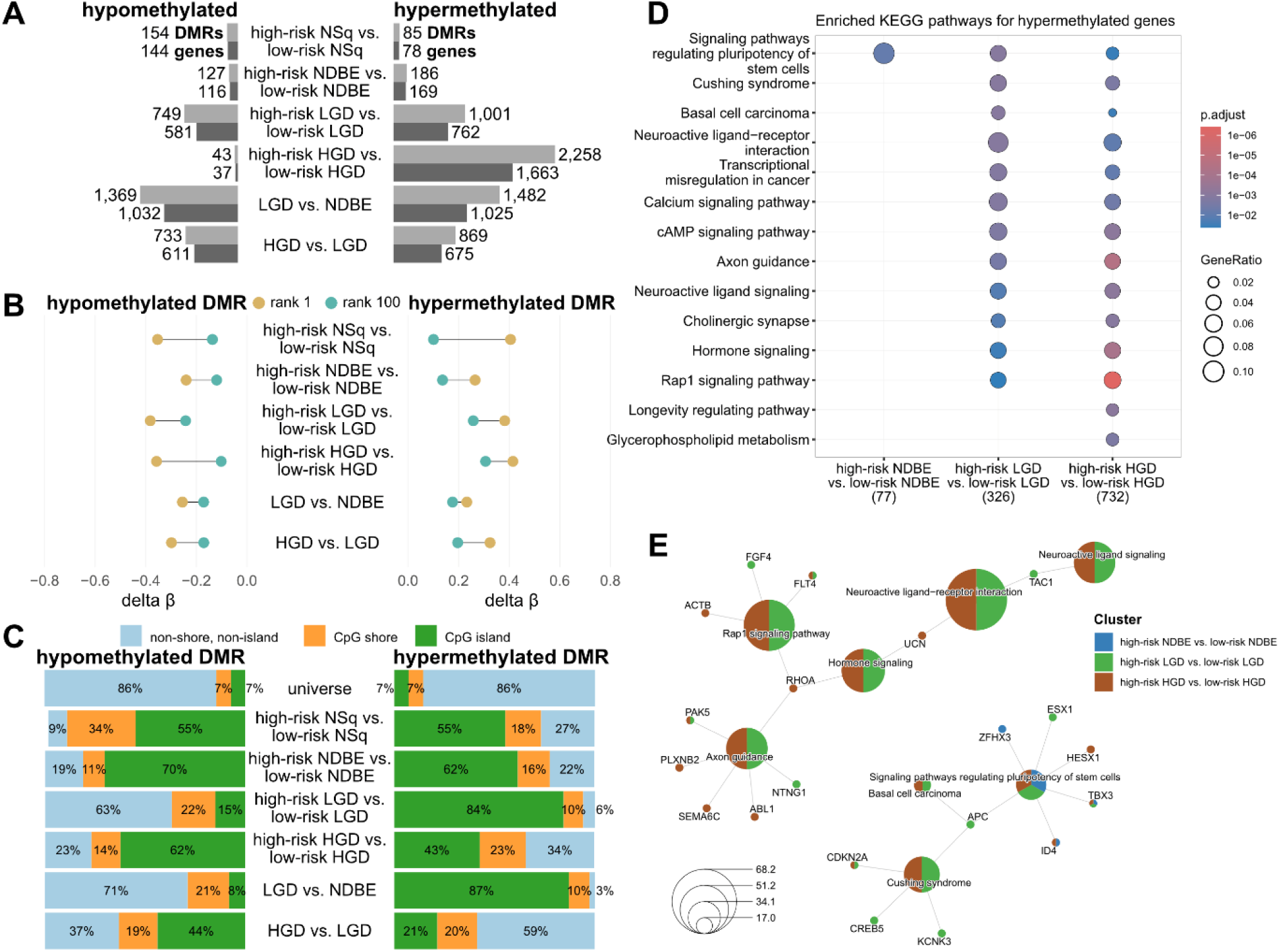
Overview of DMRs from risk-based comparisons. **(A)** Tallies of differentially methylated regions (DMRs, light gray) and genes (dark gray) across four risk-based comparisons (all EAC samples are “high-risk”, hence no valid comparator). Frequency of DMRs and genes are similar to onset/worsening of dysplasia (LGD vs. NDBE and HGD vs. LGD respectively). **(B)** Dumbbell plot showing the extent of methylation level changes of the DMRs (delta beta; rank 1 in brown, rank 100 in green). **(C)** Genomic contexts of DMRs, classified by CpG island (green), CpG shores (orange), or neither (blue). The universe refers to the genomic context of all CpG dinucleotides with valid data. **(D)** KEGG pathways enriched with hypermethylated genes across sequential EAC progression. Dysregulated pathways could occur across multiple pairwise comparisons (denoted by circles). Colors (red/blue) indicate *p* value significance; circle sizes indicate the fraction of genes in that pathway that were hypermethylated. **(E)** Gene networks highlighting the presence of known EAC and top 50 most hypermethylated genes. Genes were colored by disease classification: NDBE (blue), LGD (green), and HGD (brown).

### Differential methylation changes are independent of copy number variation

EAC is a cancer with often unstable ploidy and copy number gains within the genome (NAEINI *et al*. 2023). One major benefit of using whole methylomes over array-based methods is the ability to correlate changes in methylation with CNVs in the genome. Along the NSq-to-EAC progression pathway, there were no consistent large-scale copy number changes across all our samples (Supplementary Fig. 3). Likewise, there were few focal amplifications (0.06%) and deletions (0.21%) present across > 30% samples (20 kb windows; Supplementary Fig. 4; Supplementary Table 23). Samples further along the disease progression spectrum were, in general, associated with more extensive copy number changes across their genome, but there were no consistent CNVs that could be associated with particular disease classification (Supplementary Figs. 3, 4). We aggregated the methylation levels of CpGs by copy number variation type. While deletions were associated with a concomitant increase in methylation levels (and vice versa), the effect size of the change is small (β ± 0.01– 0.02; Supplementary Fig. 5).

The copy number of some tumor suppressor genes and oncogenes, e.g., deletions in CDKN2A (BIAN *et al*. 2002; ZAMANI *et al*. 2026), TP53 (FISHER *et al*. 2017; REDSTON *et al*. 2022) or amplification in ERBB2 (DAHLBERG *et al*. 2004; AUGUSTIN *et al*. 2022), MYC (SARBIA *et al*. 2001) and GATA6 (LIN *et al*. 2012; ZAMANI *et al*. 2026) respectively, have been previously reported to associate with the transition to metaplasia or dysplasia. In general, while the amplification/deletion of these cancer-associated genes were in line with previous reports, some genes had surprising departures from prior expectations (Fig. 3A). The deletion of CDKN2A, observed in 11 of 25 (44%) NDBE samples, was not present in higher proportions in worse pathologies (LGD: 33%; HGD: 36%; EAC: 40%). TP53 and ERBB2, both reported to have copy number changes at the transition to metaplasia, did not affect a single NDBE sample in this present study. GATA4/6 amplification, associated with EAC onset, were also present in NDBE samples (B18 and B32; Fig. 3A). The two genes with the most consistent changes were FHIT and WWOX; both genes are > 1 Mb long and span known fragile sites that are often deleted in multiple tumor types (ILIOPOULOS *et al*. 2006). However, CNV in both genes did not correlate with directional changes in methylation levels: across samples with deletions in either gene, there was a mix of hypermethylated and hypomethylated loci (Fig. 3A). Risk of worse adjacent pathology did not associate with the presence of CNVs or directional methylation shifts (Fig. 3A).

**Fig 3.**
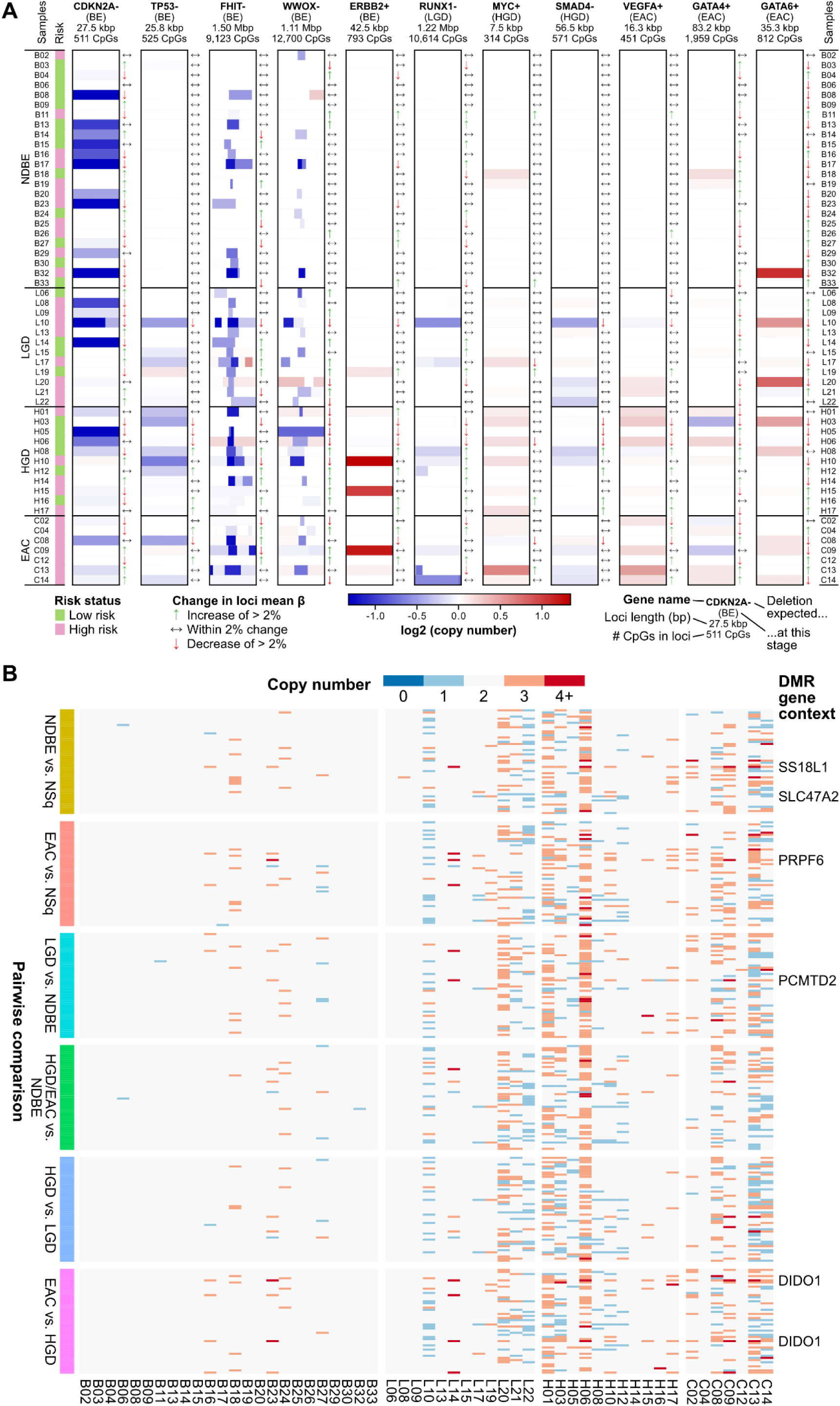
Correlation of differential methylation and copy number variation of EAC-related genes. **(A)** For tumor suppressor and oncogenes previously reported to exhibit copy number variation at different stages of EAC progression, copy number variation (blue: deletion; red: amplification) were overlaid with changes in methylation levels (β). **(B)** Top 50 hypermethylated DMRs from all six clinically relevant pairwise comparisons were overlaid with copy number information. DMRs were labelled (“DMR gene context”) if they resided in a CNV in > 20% samples. Copy number are denoted along the blue/red scale, darker hues denoting larger changes in CNV.

We investigated whether DMRs reside in genomic loci with CNVs. Focusing on the top 50 hypermethylated DMRs in every pairwise comparison, none of the DMRs resided in CNVs across > 30% samples. A more permissive consistency filter of > 20% samples reveal six DMRs (1.7%) in loci with CNVs (Fig. 3B); only one (SS18L1) of the six had previously reported association with EAC progression (MOURIKIS *et al*. 2019). None of these six hypermethylated DMRs were present in copy number reduced loci (the expected scenario; Fig. 3B). When this analysis was extended to all hypermethylated DMRs, 1.0% of all DMRs resided in CNVs across > 20% of samples; and < 0.2% across > 30% samples (data not shown). In short, the vast majority of DMRs are not associated with CNVs.

**Supplementary Fig. 3.**
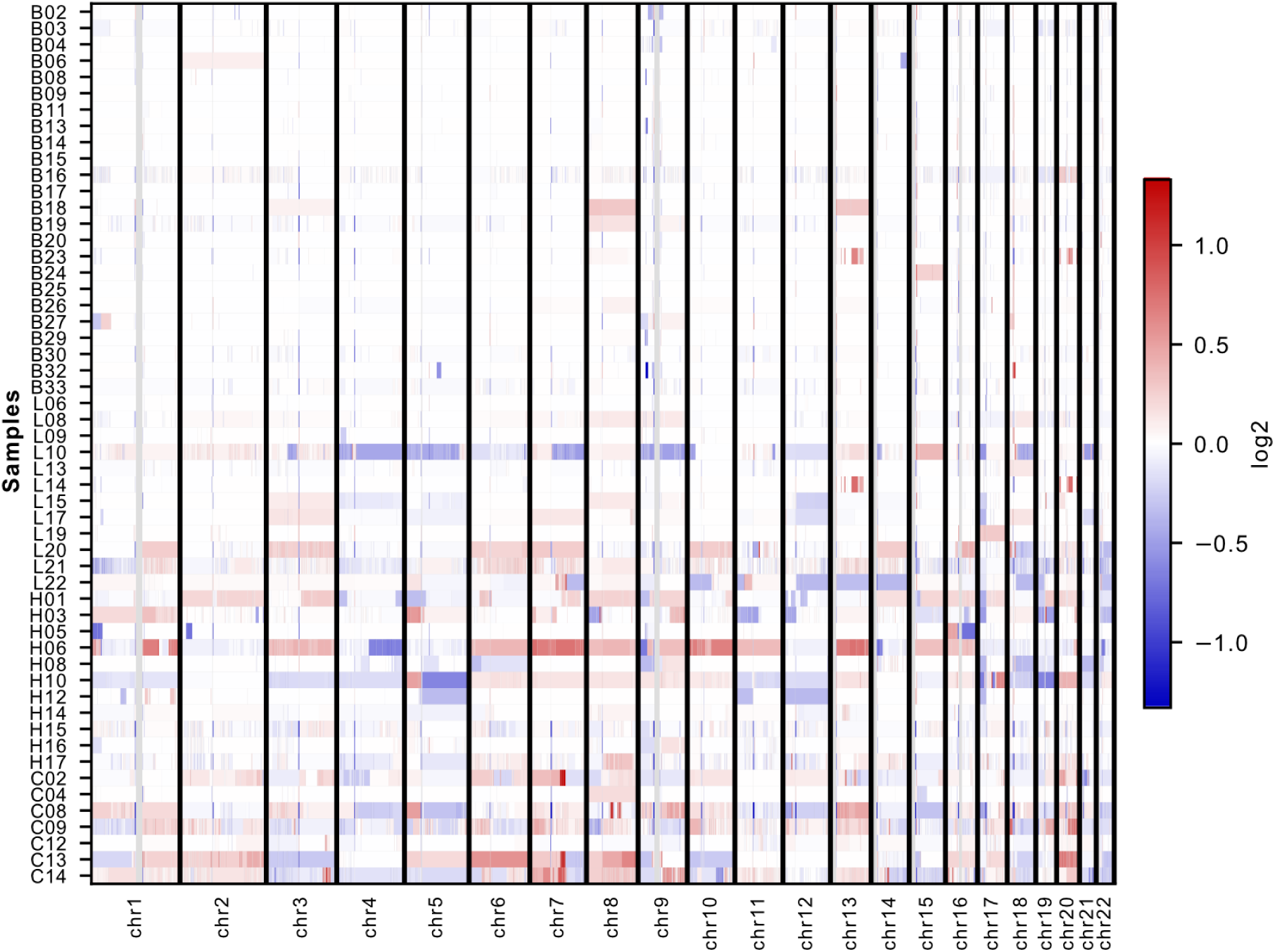
Heatmap showing CNV across all samples with sufficient coverage. There are no consistent large-scale amplifications (in red) or deletions (in blue) across samples. The vertical lines (in gray) in this plot are genomic regions known to be challenging for reads to map against and excluded from the analysis.

**Supplementary Fig. 4.**
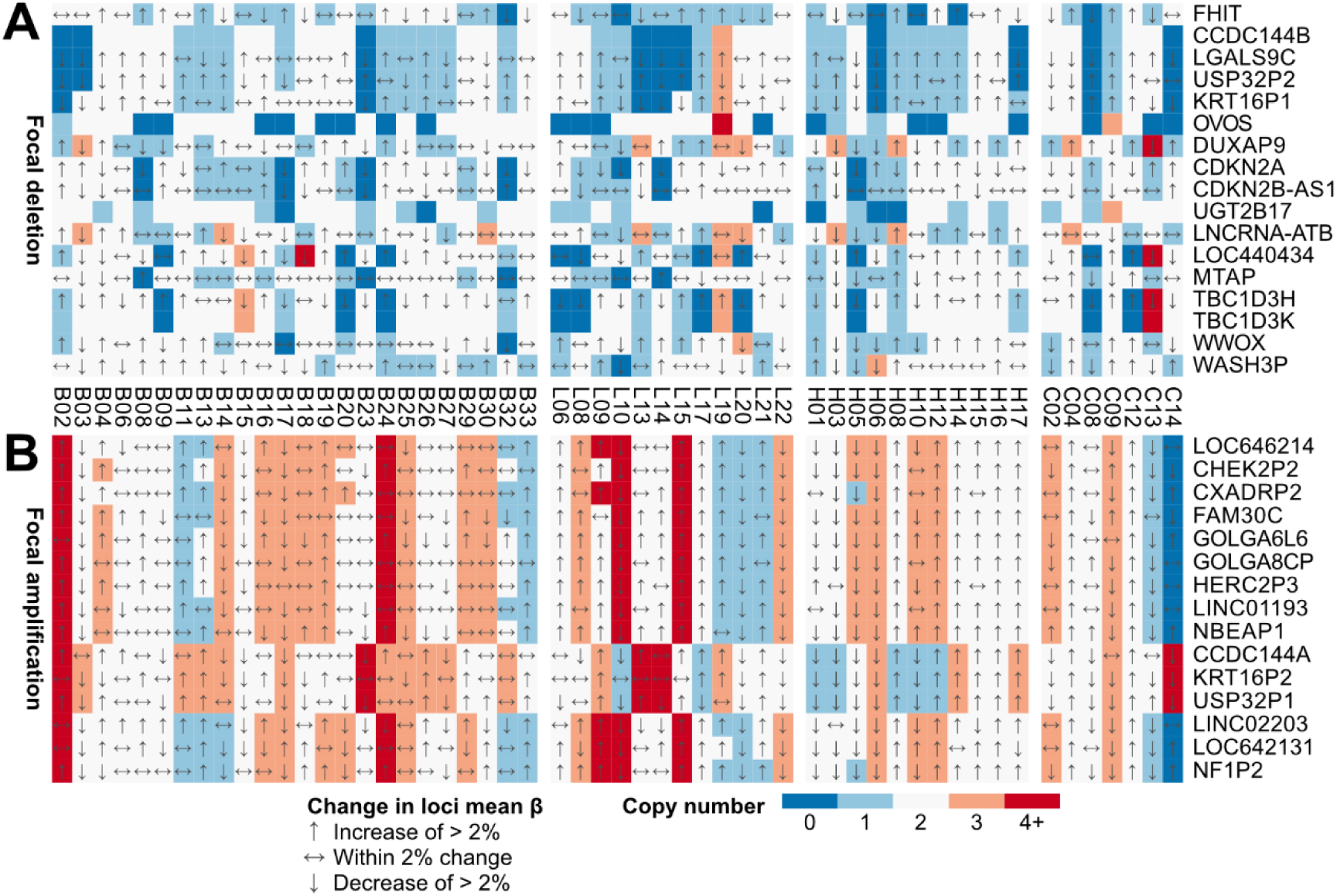
Heatmap of genes with consistent CNV in samples with sufficient coverage. Threshold for consistency was set at > 30% of included samples. Each row represents a 20 kb window contained within the annotated gene. Genes were sorted in descending order by the overall frequency of (A) focal deletions (in blue) and (B) focal amplifications (in red) across all samples; ties broken alphabetically. The mean methylation level of each 20 kb window, relative to the overall mean across samples sharing the same disease classification, are denoted by arrow direction: up for hypermethylation of > 2%, down for hypomethylation < –2%, and sideways for methylation levels within 2% of the mean.

**Supplementary Fig. 5.**
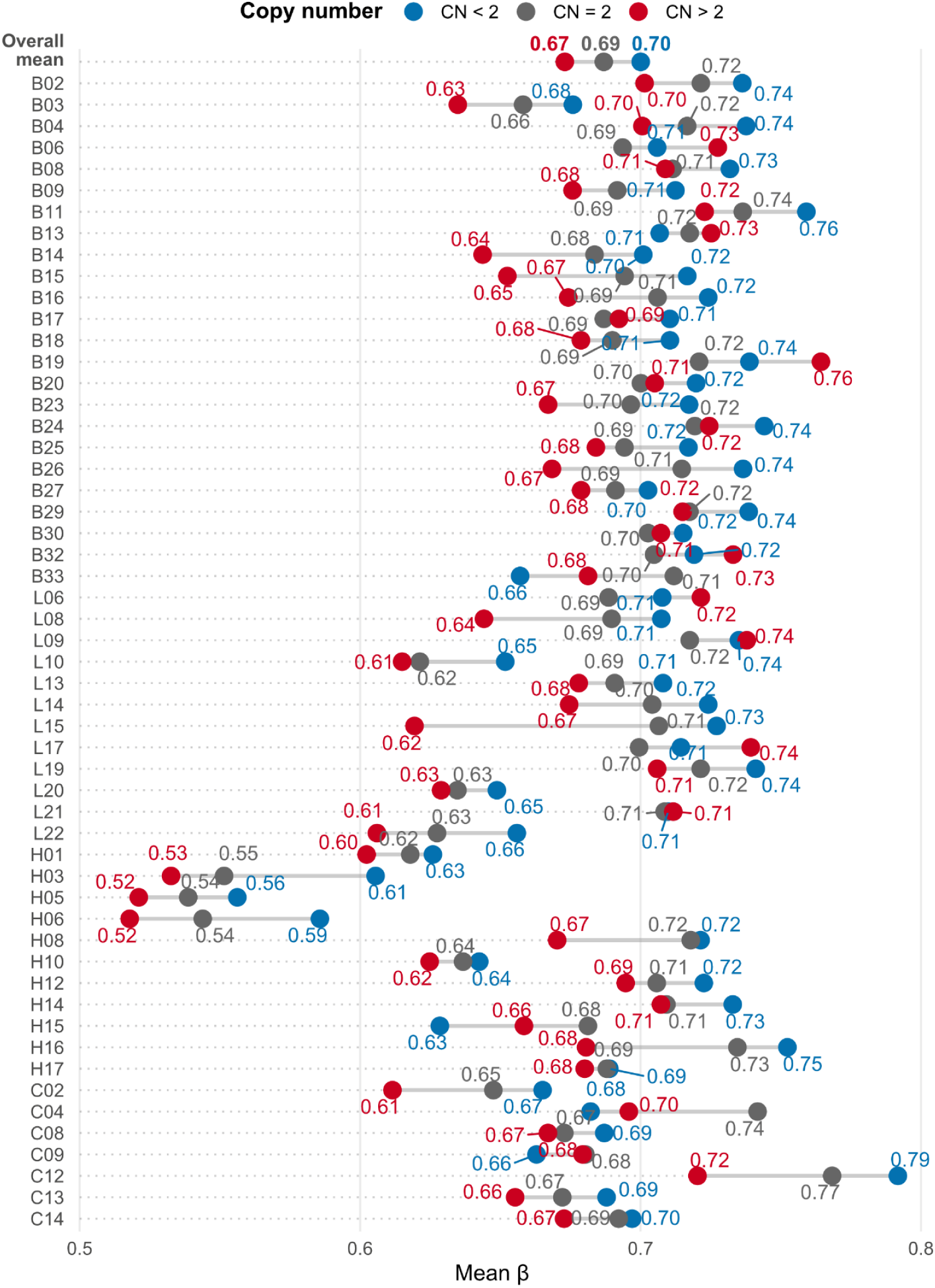
Dumbbell plots of per-sample methylation levels (mean β) across all CpGs, binned by copy number context. Averaged across all samples, genomic regions with reduced copy number (“CN < 2”; blue) had a small increase in methylation levels (β +0.01); while regions with increased copy number (“CN > 2”; red) had a similar reduction (β –0.02), relative to copy number neutral regions (“CN = 2”; gray).

### Correlation of promoter methylation with expression

As both EM-seq and RNA-seq (UDUMANNE *et al*. 2026) datasets shared a common set of *n* = 46 samples (Fig. 1A) with representation across all pathology stages, we opted to directly compare methylation against expression levels on a sample-by-sample basis, reducing the variability introduced from the use of different biological samples. While more recent studies have challenged the long-held assumption about the association of promoter hypermethylation with gene silencing (SMITH *et al*. 2020), broadly, we did observe more genes behaving in the assumed manner (i.e., anticorrelation between promoter methylation and gene expression levels; Supplementary Fig. 6).

We focused on investigating whether promoter hypermethylation correlated with gene silencing, as hypermethylated amplicons may be more useful for diagnostic tests. Significant anticorrelation between promoter methylation and gene expression was more frequent at the transition to metaplasia than other pairwise comparisons (408 NDBE vs. NSq, 11 LGD vs. NDBE, 2 HGD vs. LGD, 9 EAC vs. HGD; Supplementary Table 24). The anticorrelations were likewise stronger (more negative Pearson *r*) at the transition to metaplasia, as NSq samples tended to have lowest levels of promoter methylation and highest levels of gene expression (Supplementary Fig. 7).

**Supplementary Fig. 6.**
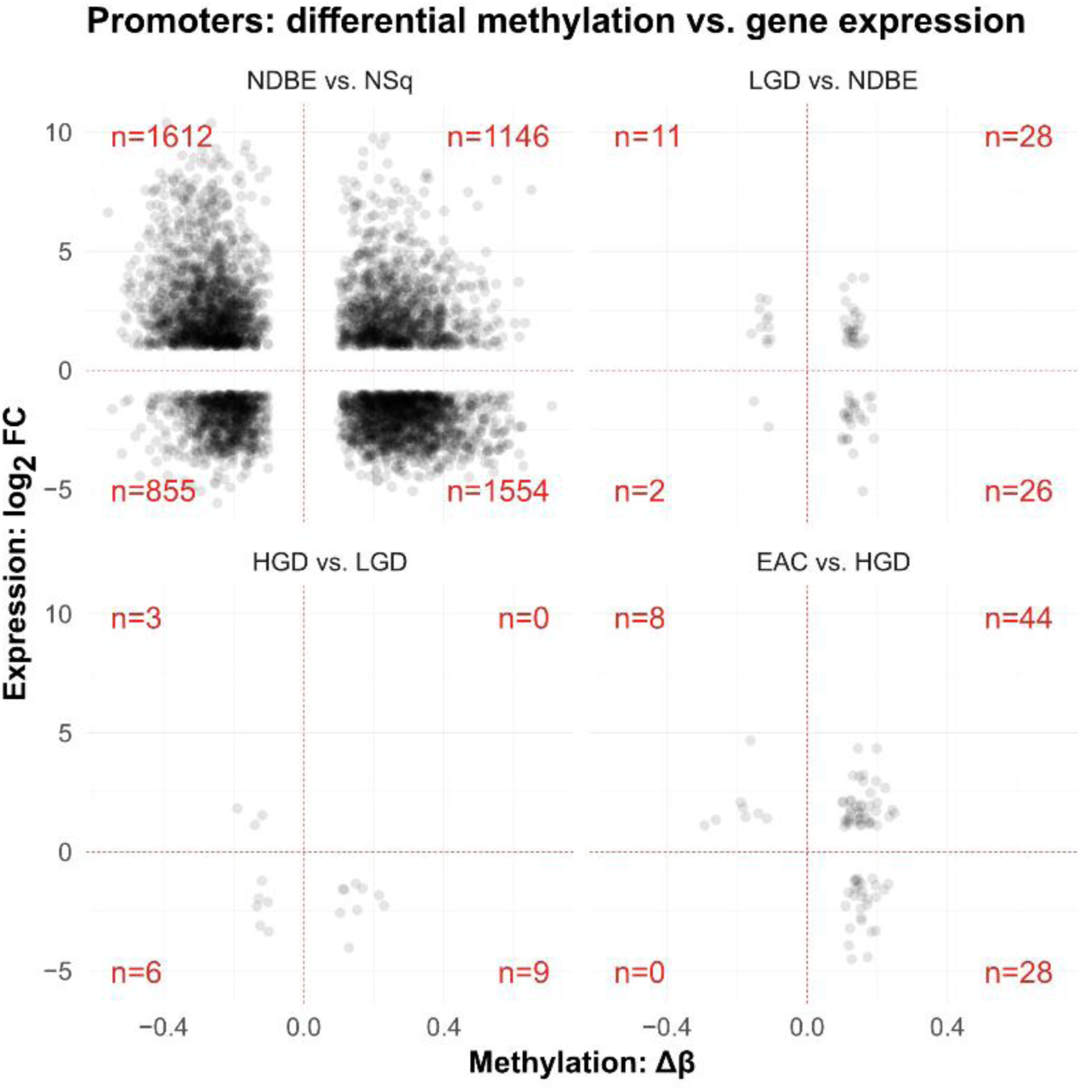
Correlation of differential promoter methylation and differential gene expression, split by pairwise comparison. Individual quadrants were tallied (red text in corners) to investigate overall trends. Genes where changes in promoter methylation and gene expression were anticorrelated would be located at either the top-left and bottom-right quadrants of each plot. Differential changes in gene expression are expressed as log2-transformed fold changes (y-axis), while differential methylation changes are untransformed differences in beta values (x-axis).

**Supplementary Fig. 7.**
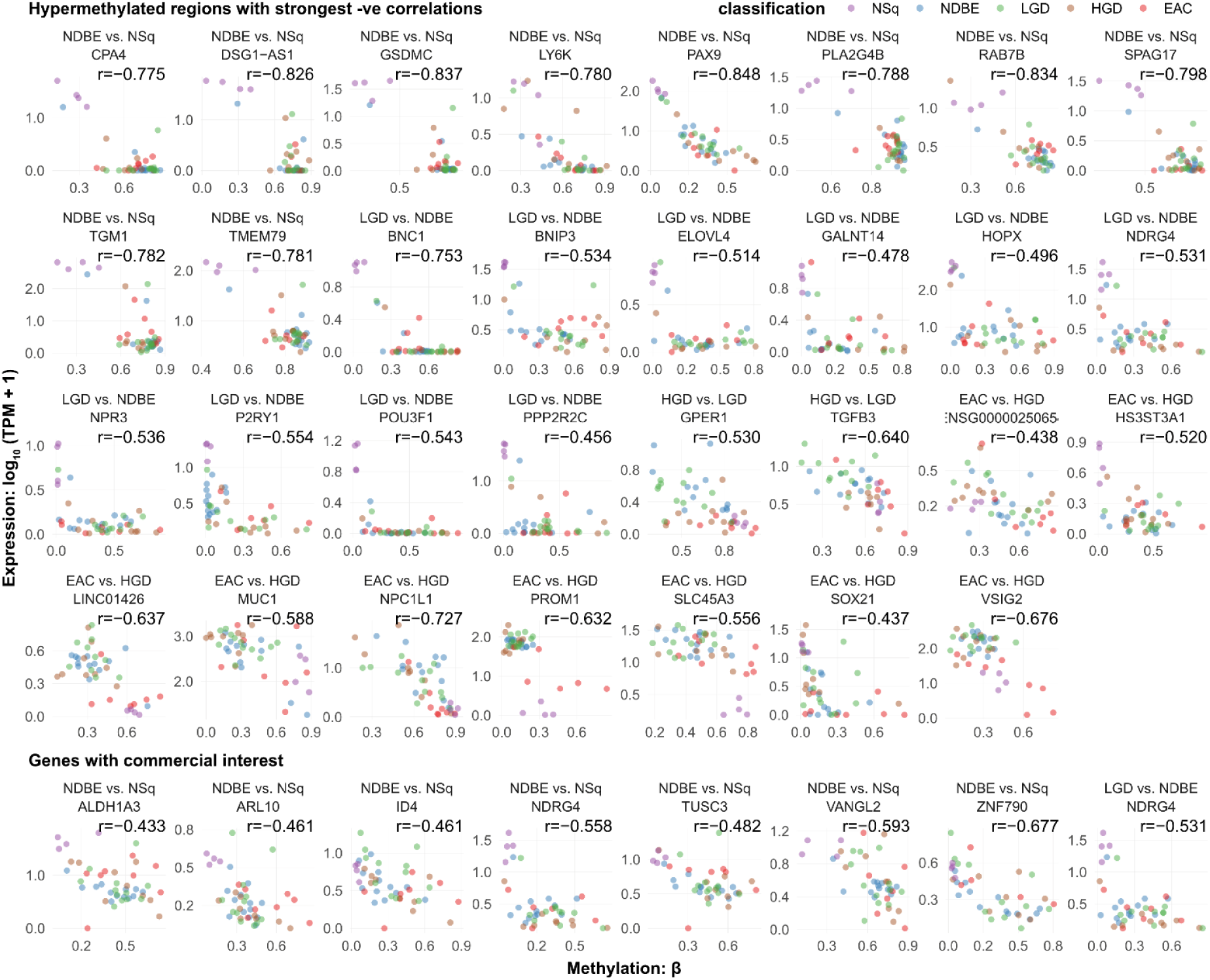
The strongest and most numerous correlations between differentially hypermethylated promoters and differentially silenced genes were at the transition to metaplasia. Each subplot represents a gene with differential changes in promoter methylation and gene expression; each point is a sample with both methylation (EM-seq) and expression (RNA-seq) data, colored by disease classification. NSq: normal squamous, purple; NDBE: non-dysplastic Barrett’s esophagus, blue; LGD: low-grade dysplasia, green; HGD: high-grade dysplasia, brown; EAC: esophageal adenocarcinoma, red.

### Deconvoluted cellular compositions reveal four distinct methylation signatures

Previous array-based studies on esophageal adenocarcinoma methylation unearthed functional signatures by sub-selecting CpGs (∼0.5%) with the greatest variability in methylation levels (XU *et al*. 2013; KRAUSE *et al*. 2016; YU *et al*. 2019; JAMMULA *et al*. 2020); or relied on expression data for initial signatures before correlating methylation patterns to said signatures (NAEINI *et al*. 2023). A similar approach failed to produce meaningful biological insights from our whole methylome data; neither did constraining the focus of the analysis on CpG islands produce anything meaningful (data not shown).

Leveraging the advancements in whole-methylome-specific deconvolution algorithms and publicly available tissue-specific methylation atlases, we used CelFiE-ISH (UNTERMAN *et al*. 2024) with the most comprehensive methylation atlas (LOYFER *et al*. 2023) to reveal four distinct methylation signatures (Fig. 4A). Loosely ordered in increasing disease severity, these signatures reflect the respective abundances in squamous epithelia (all NSq + NDBE with 20–70% squamous epithelium), columnar epithelia (majority NDBE/LGD), lymphoid tissue (mix of NDBE/LGD/HGD/EAC) and erythroid progenitors (LGD/HGD/EAC). Clusters tended to have a “binary” presence of 1–3 specific tissues, e.g., samples in the “squamous” cluster were abundant in head and neck epithelia and depleted in all other signature tissues that better defined other clusters (Supplementary Fig. 4). Risk of worse adjacent pathology did not correlate with any of the four cell type signatures (Fig. 4A).

**Fig. 4.**
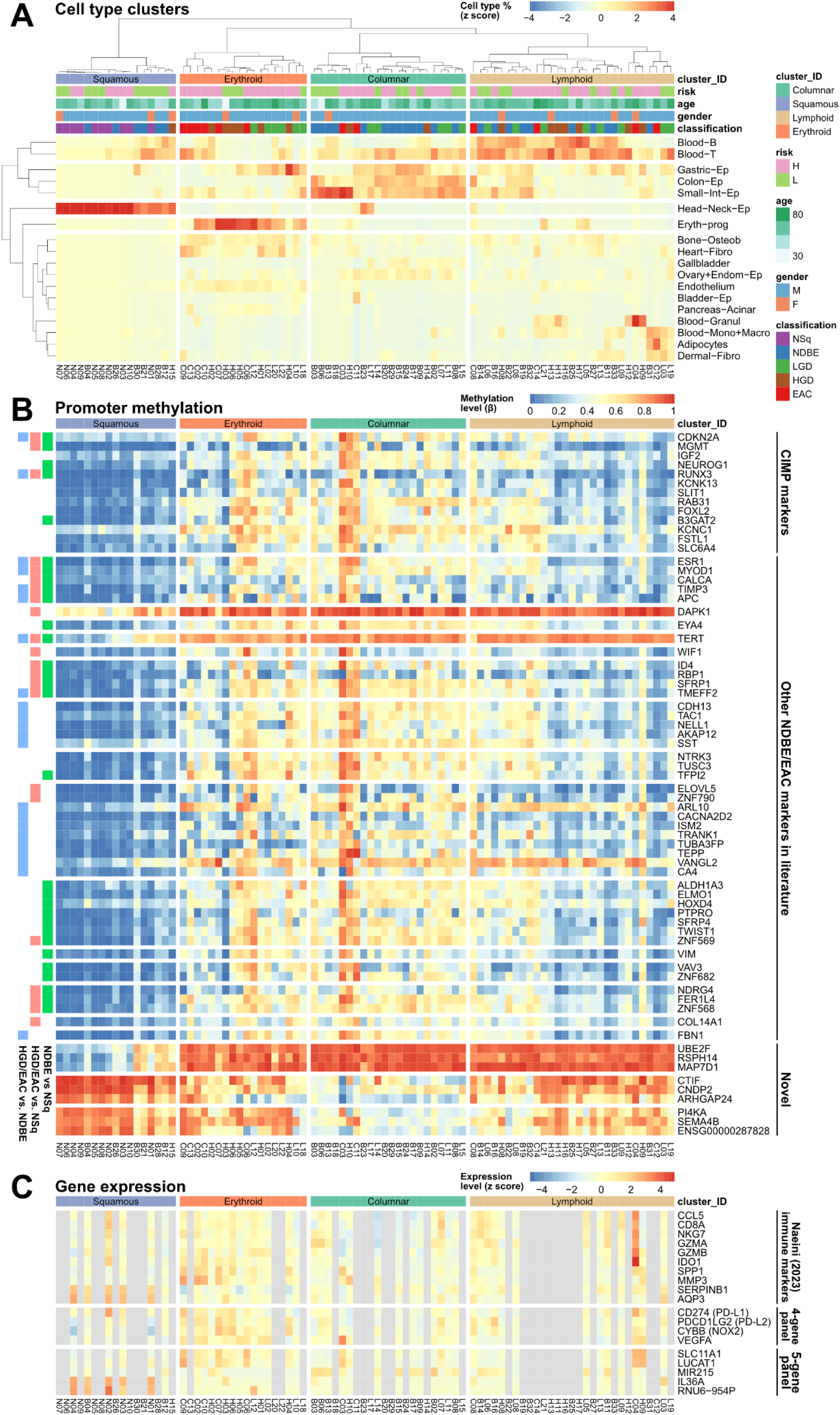
Existing methylation and expression markers perform well in distinguishing cell-type clusters derived from deconvoluted whole-methylome signatures. **(A)** Unsupervised hierarchical clustering on estimated tissue proportions reveal four clusters with differing cell-type compositions: the “squamous” cluster (abundant head and neck epithelia; blue), “columnar” (enriched in columnar gut epithelia; green), “lymphoid” (abundant lymphoid tissue; brown), and “erythroid” (high erythroid progenitor cells; orange). Heatmap values are in z-scores (normalized per tissue type; blue/red scale). **(B)** Promoter methylation for CIMP and commercially relevant markers, ordered by cell types. Markers were annotated, when possible, with its intended clinical usage (NDBE vs. NSq in green; HGD/EAC vs. NSq in red; HGD/EAC vs. NDBE in blue). Heatmap values are untransformed methylation levels (beta values; blue/red scale). **(C)** Expression patterns of proposed markers for disease progression in samples with expression data (gray: no available data). Heatmap values are in z-scores (normalized across each gene; blue/red scale).

We sought to confirm whether previous array-informed promoter methylation markers for NDBE or EAC performed well in our whole methylome dataset with two additional pathological stages. Regardless of the intended purpose (some markers were designed to separate EAC from NDBE; Supplementary Table 25), as expected, almost all markers are hypomethylated in samples high in squamous epithelia (Fig. 4B). Unfortunately, many of these markers are also hypomethylated in samples with low amounts of columnar epithelia (i.e., a negative methylation test is more likely driven by the absence of columnar than presence of squamous epithelia). Two exceptions are DAPK1 (BROCK *et al*. 2003) and TERT (CLÉMENT *et al*. 2006), as hypomethylated promoters that more reliably indicated the presence of squamous than absence of columnar epithelia. We include three novel markers (UBE2F, RSPH14 & MAP7D1) with similar properties (Fig. 4B; Supplementary Table 26), made possible by the breadth of our dataset. Markers for other signatures include three (CTIF, CNDP2 & ARHGAP24) optimized for lymphoid vs. columnar, and another three (PI4KA, SEMA4B & ENSG00000287828) for erythroid vs. columnar (Fig. 4B; Supplementary Table 26). We note however (Supplementary Table 27) that there is a weak correlation between histopathological classification (NSq/NDBE/LGD/HGD/EAC) and deconvolution cell types, apart from squamous and NSq (88.6–97.5%; Supplementary Table 27).

As there were 46 (of 86, 53.5%) samples with expression data, we investigated whether previously reported immune signatures, clustered primarily through RNA expression data (NAEINI *et al*. 2023), were similarly informative with our methylation-informed clusters. There was a strong similarity between our erythroid cluster and the previously identified high-myeloid, low-lymphocyte “immune suppressed” signature, characterized by the significant relative overexpression of SPP1/MMP3 and underexpression of SERPINB1/AQP3 (Fig. 4C; Supplementary Fig. 9). This immunosuppressed environment was further supported by the overexpression of a four-gene panel of PD-L1, PD-L2, NOX2 and VEGFA (LONG *et al*. 2022). A five-gene panel proposed recently (UDUMANNE *et al*. 2026) includes two genes (SLC11A1 & LUCAT) specific for erythroid, one (MIR215) for columnar epithelia, and two for squamous epithelia (IL36A & RNU6-954P; Fig. 4C).

**Supplementary Fig. 8.**
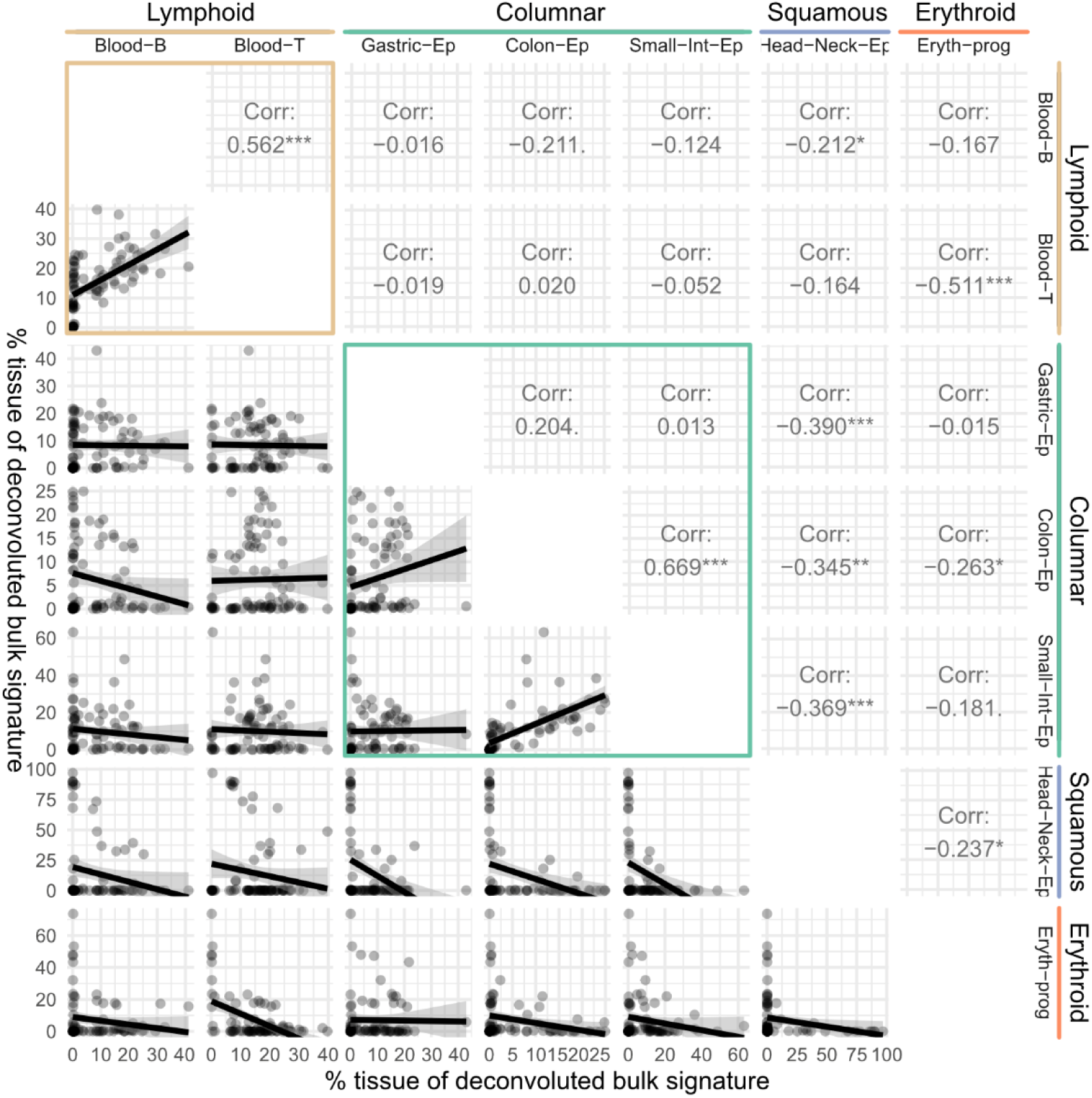
Cell-type clusters consist of a distinct set of tissues. Proportions of tissues within each cluster with more than one tissue (i.e., lymphoid, columnar) were always positively correlated intra-cluster (brown and green boxes). These proportions are almost always negatively correlated with other tissues outside of the cluster. Scatterplots with lines-of-best-fit are in the bottom triangle of the pairplot, while Pearson *r* values are in the top triangle. ***: *p* < 0.001; **: *p* < 0.01; *: *p* < 0.05.

**Supplementary Fig. 9.**
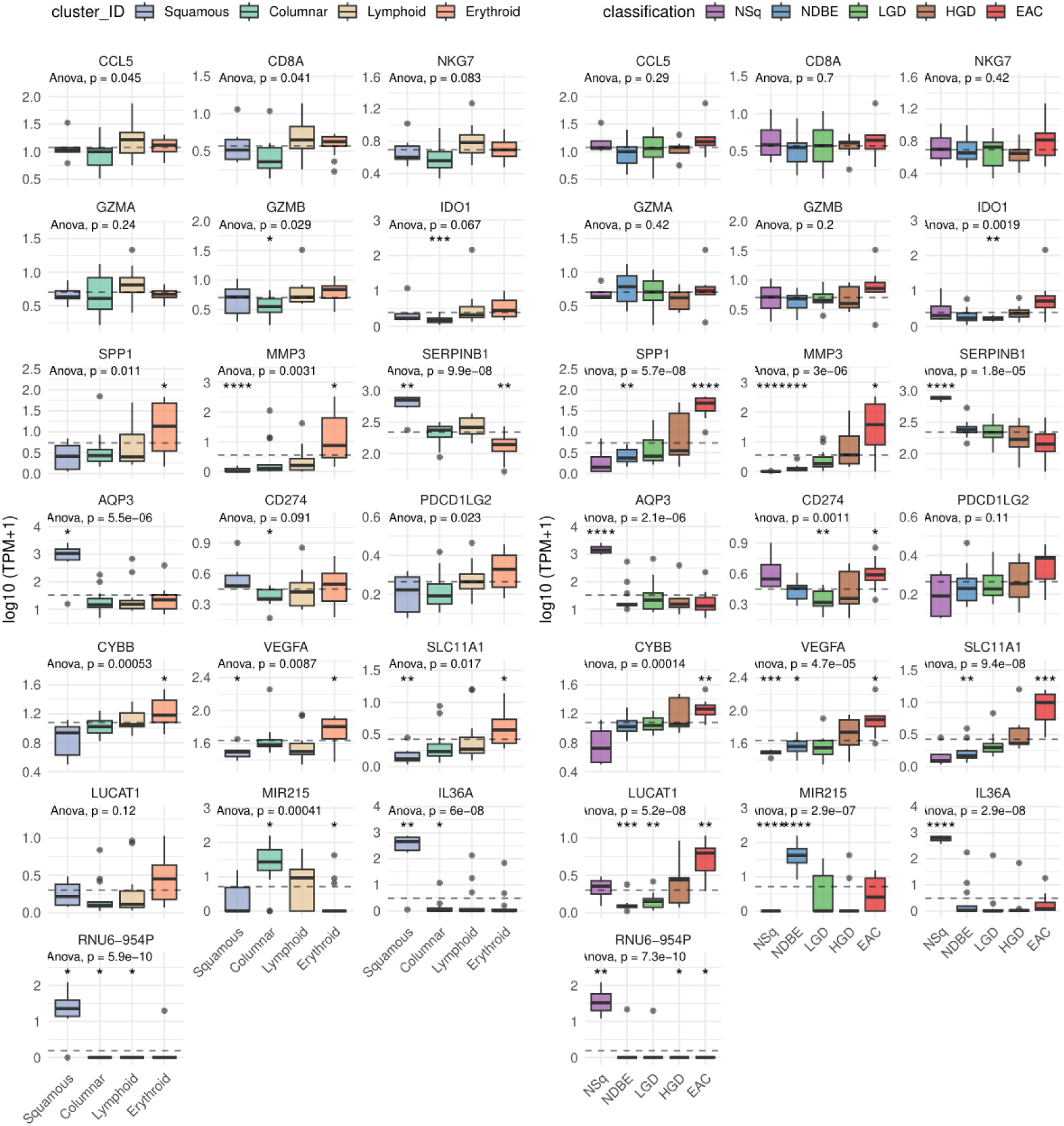
Expression of key genes in methylation-informed cell-type clusters replicate broad trends along disease progression. Per-gene expression levels (in log-transformed transcripts-per-million; y-axis) were visualized in individual box-and-whiskers plots for cell-type clusters (left panel; x-axis) and disease classification (right panel; x-axis). ANOVA *p* values denoted whether group means were at all different; asterisks denote significance of *t*-tests of each category relative to overall mean (dashed gray line). ****: p < 0.0001; ***: *p* < 0.001; **: *p* < 0.01; *: *p* < 0.05.

## Discussion

The presence of dysplasia is the main risk factor for the development of EAC in patients with BE; unfortunately, histopathological examination is unreliable for the diagnosis of dysplasia (especially LGD) (MONTGOMERY *et al*. 2001; SONWALKAR *et al*. 2010). Additional molecular evidence could assist pathologists with these difficult diagnoses, which would help guide patient management, for example regarding the decision to perform endoscopic eradication therapy (EET) for BE (RUBENSTEIN *et al*. 2024). Previous studies surveying widespread changes in the methylation landscape across the NSq-to-EAC spectrum did so by including samples primarily from three stages: NSq, NDBE and EAC (XU *et al*. 2013; KRAUSE *et al*. 2016; YU *et al*. 2019; JAMMULA *et al*. 2020; NAEINI *et al*. 2023). Building upon the long-term efforts of the PROBE-NET collaboration, we sequenced novel methylomes from the harder-to-distinguish LGD (*n* = 19) and HGD (*n* = 16) stages. With EM-seq, we also obtained additional genome-wide evidence from the three better-studied stages (NSq, NDBE, and EAC), expanding the breadth of the gathered evidence to > 25M cytosines across the genome from the previously available < 1M data.

With the expanded breadth, we observe that EAC progression involves global hypomethylation (Fig. 1C) and focal hypermethylation (Fig. 1E), similar to esophageal squamous cell carcinoma (HOSHIMOTO *et al*. 2015) and other cancers (NISHIYAMA & NAKANISHI 2021). In line with previous reports, global hypomethylation occurred mostly in gene bodies and intergenic regions, with focal hypermethylation centered on CpG islands (Fig. 1E). Echoing the “Anna Karenina principle” (also proposed for microbiomes (ZANEVELD *et al*. 2017), whereby healthy microbiomes share greater overlap than dysbiotic ones), overall methylation patterns were the most similar among NSq samples, becoming increasingly dissimilar and dysregulated along the disease progression spectrum (Fig. 1D). These changes were not gradual—rather, punctuated—at two specific stages. Both in terms of frequency (Fig. 2B) and magnitude (Fig. 2C), the largest differential methylation changes occur at the transition to metaplasia (NDBE vs. NSq), followed by the onset of invasive cancer (EAC vs. HGD); the same stages are considered less difficult histopathological diagnoses, compared to dysplasia. This is similar to prior efforts in which the top 5000 most variable methylation probes differentiated NSq well from NDBE and EAC, but not EAC from NDBE (KRAUSE *et al*. 2016); and similarly, the most extensive transcriptomic changes occur at the transition to metaplasia (UDUMANNE *et al*. 2026). While there were DMRs that could close the diagnostic gap in distinguishing the onset of dysplasia (LGD vs. NDBE) and worsening dysplasia (HGD vs. LGD), they were roughly an order fewer in number and half the magnitude, paralleling the difficulties faced in histopathology. Hypermethylated DMRs were disproportionately located within CpG islands (Fig. 2D), as expected, and were in genes modulating cell growth and cell-cell adhesion (Figs. 2E, 2F).

The more even spread of coverage from the use of enzymatic conversion (VAISVILA *et al*. 2021; GUANZON *et al*. 2024) enabled the more accurate identification and analysis of CNV with disease progression. We did not seek to replicate the finding of widespread genomic alterations in HGD/EAC samples well established previously (NONES *et al*. 2014; ROSS-INNES *et al*. 2015; SECRIER *et al*. 2016; NAEINI *et al*. 2023; ZAMANI *et al*. 2026); rather, we sought to understand whether methylation levels and DMR identification in EAC were confounded by CNV, as with other cancers (MARTIN-TRUJILLO *et al*. 2017; SUN *et al*. 2018). Overall, while we did observe a slight increase of methylation in deleted regions, and vice versa in amplified regions, the absolute magnitude (1–2%; Supplementary Fig. 5) is minor compared to the typical deltas for biomarker amplicon design (10–40%). The methylation levels of genes with known CNV did not shift in a meaningfully consistent manner (Fig. 3A), and few identified DMRs were in loci with CNV (Fig. 3B). Our data suggest that for EAC, the changes in methylation were mostly independent of CNV. Hence, the use of CNV-blind methods to assay methylation (e.g., microarrays, qPCR/dPCR) remains appropriate for biomarker discovery and diagnostics development.

The predominant association of promoter hypermethylation with gene silencing has been previously documented in NDBE/EAC (KRAUSE *et al*. 2016; JAMMULA *et al*. 2020); we assessed whether similar trends held in the dysplastic stages (LGD/HGD) novel to this study. Focusing on the *n* = 46 samples with both transcriptomic and methylome data, in line with expectations, promoter methylation levels were mostly anticorrelated with gene expression in all four pairwise comparisons across the disease spectrum (Supplementary Fig. 6). Differential promoter methylation and gene expression were the strongest and most numerous from the transition to metaplasia (NDBE vs. NSq; Supplementary Figs. 6, 7), underscoring the difficulty in developing diagnostics for the progression to dysplasia or carcinogenesis.

The prioritization of methylation markers with additional multi-omic evidence (RNA expression) has recently been published for esophageal squamous cell carcinoma (XI *et al*. 2022). In other cancers, some commercially available assays of promoter hypermethylation are known to strongly associate with the silencing of the same gene, e.g., MGMT for glioblastoma (LOCKE *et al*. 2019) and GSTP1 for prostate cancer (PIDSLEY *et al*. 2024), influencing clinical management of those diseases (HEGI *et al*. 2005; MALDONADO *et al*. 2014). Our *in silico* testing of commercially available NDBE/EAC biomarkers on our samples revealed the standout marker NDRG4 (IYER *et al*. 2020). As well as having strong anticorrelation with expression, the promoter hypermethylation not only satisfied its original design intention of detecting the transition to metaplasia (NDBE vs. NSq); it also performed well in detecting onset of dysplasia (LGD vs. NDBE; Supplementary Fig. 7). We propose other potential genes that could better separate other stages, e.g., METTL7A and VSIG2 for EAC vs. HGD, as the increased methylation and reduced expression occurs at the onset of invasive cancer development (Supplementary Fig. 7).

Recent optimization of tissue-of-origin deconvolution algorithms on sequencing data and publicly available tissue-specific methylation atlases have enabled the more accurate decomposition of bulk methylome signals into their constituent tissues (CAGGIANO *et al*. 2021; LOYFER *et al*. 2023; KEUKELEIRE *et al*. 2023; UNTERMAN *et al*. 2024), as the atlas is calibrated on a wider range of tissue-specific CpGs not present on existing commercial arrays. Samples were grouped into four cell-type clusters based on their relative proportions of different tissues, somewhat consistent with BE/EAC progression (Fig. 4A). The “squamous” group, enriched with squamous epithelia, consisted of all NSq and several NDBE samples that included 20–70% squamous epithelium; the “columnar” group, enriched with columnar epithelia, was mostly composed of NDBE samples and LGD. This is followed by two immune-related groupings: “lymphoid”, enriched in B– and T-lymphocytes, with constituent samples spanning the widest range of NDBE to EAC; and “erythroid”, enriched in erythroid progenitor cells normally found in bone marrow, solely from dysplastic samples (LGD to EAC). It is important to note that the deconvolution algorithm can only assign relative abundances to tissues present in the underlying atlas, and there was no “myeloid” grouping in the atlas. Regardless, the composition of immune cells in the esophageal tumor microenvironment has prognostic implications (CUI *et al*. 2021; DAVERN *et al*. 2021).

Recent transcriptional work on EAC has revealed the correlation of patient survival with four distinct immune signatures (NAEINI *et al*. 2023). Two of their expression-derived signatures correlate well with our methylome-derived cell-type clusters. Our lymphoid cluster, high in lymphocytes and low in erythroid progenitors (Fig. 4A), matches the previously described “immune moderate” signature; while our erythroid cluster, high in erythroid progenitors and low in lymphocytes (Fig. 4A), matches the “immune suppressed” signature. This suggests that BE with dysplasia and EAC may develop in an immune-suppressed environment, compared to non-dysplastic BE. Although erythroid progenitor cells would be unexpected in BE/EAC, they are reported to have an immunosuppressive effect on T-cells (GRZYWA *et al*. 2021; BOZORGMEHR *et al*. 2023). When we overlaid transcriptional data available from a subset of samples (UDUMANNE *et al*. 2026), the transcriptional markers for “immune suppressed” from Naeini et al. (2023), i.e., high SPP1/MMP3 and low SERPINB1/AQP3, and overexpression of PD-L1, PD-L2, NOX2 and VEGFA (LONG *et al*. 2022), were similarly observed in our erythroid cluster (Fig. 4C; Supplementary Fig. 9). In short, our methylome-driven deconvolution of bulk tissues supports previous transcriptional approaches. Further insights into the immune composition of EAC tumor microenvironment hold potential therapeutic significance (LONIE *et al*. 2023).

In conclusion, we obtained methylomes with greater breadth, across all CpGs in the genome, and across a greater range, as we included two dysplastic stages with very limited prior methylation data (LGD/HGD). Overall, disease progression involves global hypomethylation with focal hypermethylation at CpG islands, with the bulk of changes occurring at the transition to metaplasia. Designing markers to address clinical gaps, i.e., accurately denoting the onset and/or worsening of dysplasia, and sampling error, remains challenging. We do acknowledge limitations in our marker discovery: power would improve with more samples, and our “low-risk” category could be misclassified due to not detecting worse adjacent pathology. Another important limitation is that the erythroid grouping indicates an immunosuppressed environment with characteristics similar to EPC expression pattern, and not necessarily that EPCs are present, due to the absence of a “myeloid” grouping in the atlas. A silver lining is that methylation levels across EAC stages are mostly independent of copy number variation, i.e., copy-number-blind methods will likely function equally well. The deconvolution of bulk methylome data into constituent tissue-of-origin has enabled unexpected results with possible insights into BE/EAC biology. We provide additional molecular evidence to improve the identification of dysplasia, with the hopes of bridging the current clinical gap.

## Supporting information

## Acknowledgements

We thank the patients who generously provided their samples for research; and the Australian Genome Research Facility (Melbourne, Australia) for library creation and sequencing.

## Author contributions

R.V.L. and J.R. conceived and designed the study. M.J.B. and R.V.L. enrolled patients. T.P.U., K.K.M.L. and R.V.L. performed sample selection. T.P.U., K.C., A.A. and K.K.M.L., carried out experiments. T.Y., P.K., D.M. and I.B. conducted the pathology assessments. Y.J.L., T.P.U., D.P., S.J.L., R.V.L. and J.R. carried out data analysis and/or interpretation. Y.J.L. wrote the initial manuscript. Y.J.L., T.P.U., D.P., S.J.L., R.V.L. and J.R. reviewed and edited the manuscript. All authors provided feedback and approved the final version of the manuscript.

## Data availability

Raw sequencing data in FASTQ format is available at SRA, with accession PRJNA1540235. Intermediate tab-delimited files with per-CpG beta values for all samples, genome used for mapping, and all associated annotation files are available at https://data.csiro.au/collection/78585. Code for analyses and figure generation is available at https://github.com/lyijin/eac_emseq.

